# A multiregional image–text dataset and benchmark for vision-language modeling of plant diseases

**DOI:** 10.64898/2026.07.01.735881

**Authors:** Trang V. Nguyen, Khang Nguyen Quoc, David Harwath, Luyl-Da Quach, Phuong D. Dao

## Abstract

Plant diseases remain a major challenge to global food production, and timely, accurate, and scalable detection of plant stress is critical to reducing these losses. Recent advances in digital imaging and artificial intelligence offer unprecedented opportunities for precision crop disease detection and management. Yet, existing plant disease datasets remain often fragmented across crop and disease systems, and are largely dominated by controlled-environment imagery. The lack of standardized, interoperable, and representative datasets limits reproducibility, transferability, and scalability of AI systems, thereby constraining their deployment in operational agricultural applications. Here we present LeafMD, an integrated multimodal plant disease dataset and benchmark resource that includes LeafNet 2.0, a large-scale multimodal digital image dataset comprising 255,855 image–text pairs across 37 crop species, 197 crop–disease classes, and 9 geographic regions spanning tropical, subtropical, and temperate agricultural systems. Unlike conventional datasets, LeafNet 2.0 integrates biologically grounded symptom descriptions with image-level annotations of early and late disease stages, enabling symptom-aware analysis of disease progression under realistic field conditions. We further introduce LeafBench 2.0 as part of LeafMD, a visual-question answering benchmark covering nine fine-grained plant pathology tasks, including pathogen classification, lesion characterization, symptom interpretation, and disease severity assessment. Evaluation across 16 vision–language models revealed substantial performance gaps between coarse disease recognition and fine-grained pathological reasoning, while agriculture-adapted models consistently outperformed several larger general-domain architectures on symptom-oriented tasks. Together, LeafNet 2.0 and LeafBench 2.0 establish LeafMD as a multimodal resource for developing disease-aware agricultural foundation models and studying fine-grained pathological reasoning in real-world environments.

## Background & Summary

Plant diseases impose substantial constraints on global food production, resulting in significant yield and economic losses across major staple crops, including maize, rice, wheat, soybean, and potato. Pathogen-induced yield reductions can reach up to 40% under severe outbreaks, contributing to an estimated USD 220 billion in annual global losses (Savary et al., 2019; Duhan et al., 2025). Across crop–pathogen systems, disease manifests on foliage as a continuum of visually distinguishable symptoms, from early discoloration and lesions to late-stage necrosis and tissue deformation, corresponding to physiological disruption during disease progression. Accordingly, leaf-level observations are widely used in plant disease detection and characterization studies (Lv et al., 2023).

Traditionally, plant disease assessment has been carried out through visual inspection by domain experts. While effective, this approach is labor-intensive and difficult to scale across large and diverse agricultural systems. Recent advances in computer vision, particularly through machine learning (ML) and deep learning (DL) methods, have enabled more automated analysis of leaf disease images and have increased the potential for large-scale disease monitoring (Ali et al., 2024; Bhatti et al., 2024; Alhwaiti et al., 2025; Ashurov et al., 2025; Hari & Singh, 2025; Bilal et al., 2025; Abdalla et al., 2025). Despite these advances, deep learning–based methods still rely heavily on the availability of large, well-annotated training datasets (Xu et al., 2024; Argüeso et al., 2020). Obtaining such datasets remains challenging due to the complexity of real-world agricultural conditions and the high cost of data collection and labeling (Sharma et al., 2022). As a result, although many public plant disease datasets have been released, limitations related to data scale, coverage, and annotation quality persist.

A common limitation of existing plant disease datasets is their limited coverage of crop species and geographic regions. As a result, many studies depend on self-collected datasets drawn from a small number of crops or locations, reflecting the limited availability of large and standardized public resources (Salka et al., 2025). This restricted scope is problematic because leaf morphology and visual appearance vary widely across crops, and disease symptoms are further influenced by regional differences in climate, soil conditions, and cultivation practices (Song et al., 2023). As a result, models trained on such datasets often perform poorly when applied to new crops or regions outside the original collection context (Li et al., 2025), limiting model’s transferability and scalability.

Beyond limited coverage, most plant disease datasets contain little semantic information. Annotations are typically provided as categorical disease labels designed for image classification or detection, which emphasizes on visual appearance but offers limited detail on symptom characteristics, disease progression, or observation context (Liu et al., 2024). Some recent datasets have extended disease representation by incorporating additional modalities, such as sensor data, temporal image sequences, and expert annotations (Xiong et al., 2025; El Sakka et al., 2025; Yuan et al., 2025).

Recent advances in large language models and multimodal foundation models, particularly systems such as GPT-4o, have created new opportunities to leverage previously underutilized plant pathology data modalities. These include disease scouting notes, extension pathology bulletins, expert diagnostic reports, fungicide trial records, and symptom descriptions embedded in historical crop disease surveys, which have traditionally remained disconnected from machine learning pipelines. In plant pathology, these models enable the transformation of heterogeneous unstructured text into biologically grounded semantic image-level captions annotations. As a result, plant disease datasets can evolve from closed-set image classification resources into multimodal knowledge repositories that support advanced tasks for current precision agriculture, such as fine-grained symptom reasoning, visual question answering, image–text retrieval, disease progression modeling, and zero-shot recognition of emerging crop–pathogen combinations.

However, existing caption-based datasets often use limited domain-specific agricultural terminology and rarely capture symptom changes over time, resulting in an incomplete representation of disease context (Xiong et al., 2025). Most existing plant disease datasets focus on disease presence or early symptom detection, with limited coverage of later stages of disease development (Demilie, 2024). In field conditions, disease symptoms evolve over time, leading to clear changes in lesion morphology, affected tissue area, and spatial distribution. Datasets that emphasize early-stage symptoms therefore provide an incomplete view of disease expression and restrict analyses related to disease staging and severity assessment.

To address the limited geographic diversity and sparse biological annotation of existing plant disease datasets, we developed LeafMD, an integrated multimodal plant disease dataset and benchmark resource for plant pathology vision–language research. LeafMD includes LeafNet 2.0, a multiregional image–caption dataset containing 255,855 images spanning 37 crop species, 144 disease classes, and 9 countries across tropical, subtropical, and temperate production systems. Rather than relying solely on categorical disease labels, LeafNet 2.0 pairs each image with symptom-focused descriptions derived from plant pathology references together with disease stage annotations, a clinically and agronomically important dimension that has remained largely overlooked in existing plant disease datasets. This design allows models to learn not only disease identity, but also how symptoms emerge and evolve under heterogeneous field conditions. In parallel, LeafMD includes LeafBench 2.0, a benchmark for evaluating fine-grained disease understanding in vision–language models. Together, LeafNet 2.0 and LeafBench 2.0 establish LeafMD as a multimodal resource for developing transferrable and scalable disease-aware agricultural foundation models and studying fine-grained pathological reasoning in real-world environments.

Our contributions are threefold:

i) We substantially expand the geographic and environmental coverage of existing plant disease resources by constructing, to our knowledge, the largest multiregional plant disease image dataset currently available.
ii) We introduce a large-scale plant disease image–caption resource in which biologically grounded symptom descriptions and disease stage information are aligned with individual images, allowing multimodal learning of symptom morphology and disease progression.
iii) We develop LeafBench 2.0, a pathology-oriented visual question answering benchmark designed to evaluate disease understanding beyond coarse recognition, including symptom interpretation, lesion characterization, pathogen identification, and disease stage reasoning under realistic agricultural conditions.

## Method

The dataset was constructed using a three-stage, sequential workflow comprising (1) image dataset collection, (2) caption construction, and (3) validation. The overall pipeline is illustrated in **Fig. 1**, and each stage is described in detail in the following subsections.

**Figure 1.**
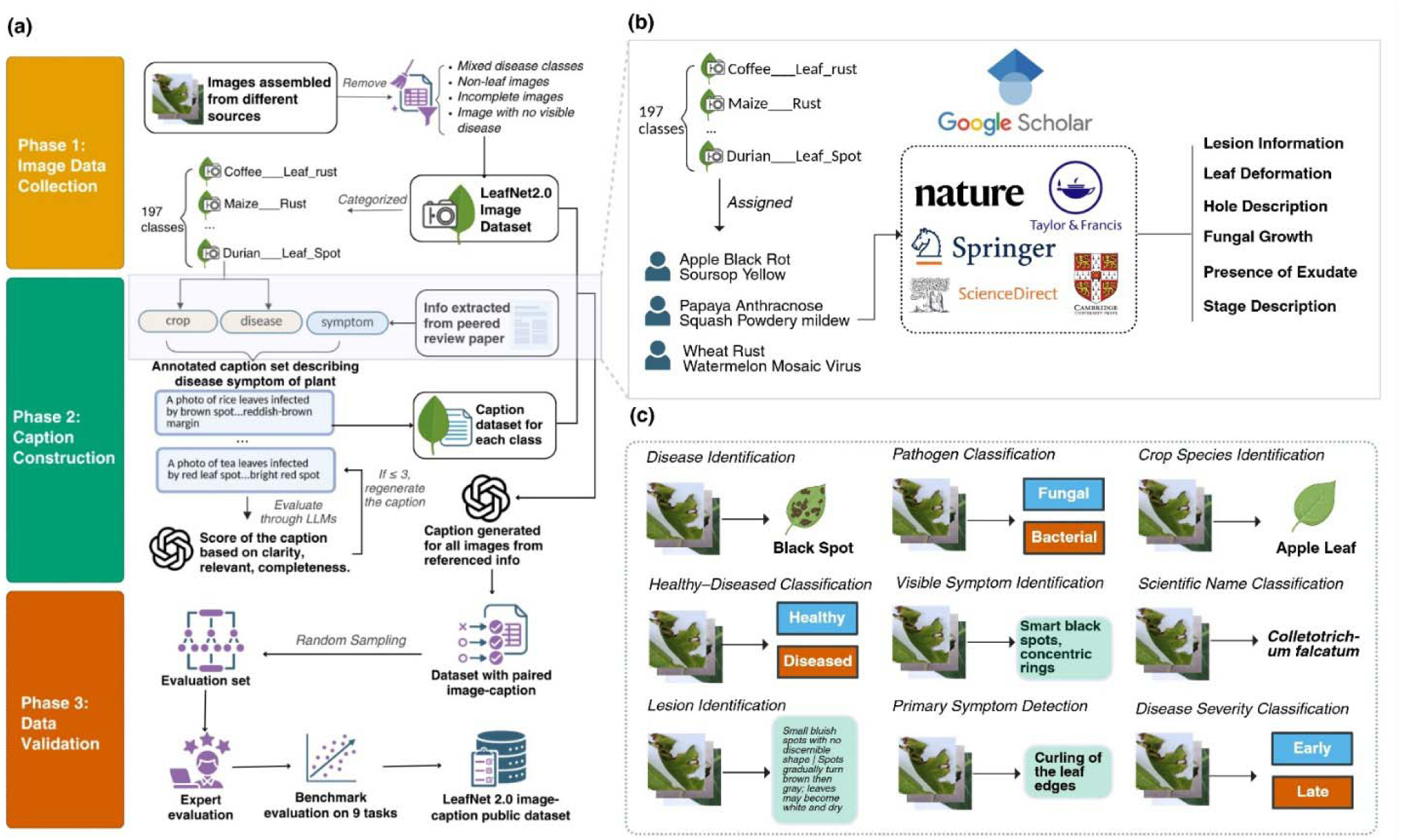
Overview of the LeafNet2.0 dataset construction process. (a) Overall workflow of the LeafNet2.0 dataset construction process, from image collection and curation to caption generation, quality evaluation, and benchmark preparation. (b) Scientific literature sources and disease-related information used to guide standardized caption construction in Phase 2. (c) Representative benchmark tasks in Phase 3 illustrating the multi-task evaluation framework supported by LeafNet2.0 for plant disease understanding.

### A. Image Dataset Collection

The image dataset was developed by expanding the original LeafNet collection (Nguyen Quoc et al., 2026), which was previously concentrated in subtropical and temperate systems (Africa and the United States), with data tropical cropping environments from South and Southeast Asia, where disease expression and field conditions differ markedly from those represented in the original collection.

Most images were compiled and standardized from publicly available plant disease datasets distributed through established data repositories and peer-reviewed data journals (e.g., Data in Brief, Mendeley Data). For datasets not publicly accessible, images were obtained through direct communication with the originating institutions following permission approval.

A total of 89,398 additional images with geographic metadata were incorporated into LeafNet2.0, increasing the dataset to 278,131 labeled samples spanning nine geographic regions, together with a small subset of images with unknown origin (1.4% of the dataset; Fig. 2a–c and Supplementary Data S1). Most samples were acquired using RGB imaging devices, primarily digital cameras and mobile phones, whereas 5.13% of the dataset lacked acquisition metadata because these details were not reported in the original sources (Fig. 2d). Compared with the original dataset, the extended collection substantially increases the proportion of images captured under natural field conditions (Fig. 2e), characterized by heterogeneous backgrounds, variable illumination, and unconstrained imaging environments. This expanded diversity in geographic coverage, acquisition devices, and environmental settings reduces the dominance of controlled laboratory-style imagery commonly observed in existing plant disease datasets, thereby providing a broader representation of real-world agricultural conditions.

**Fig. 2.**
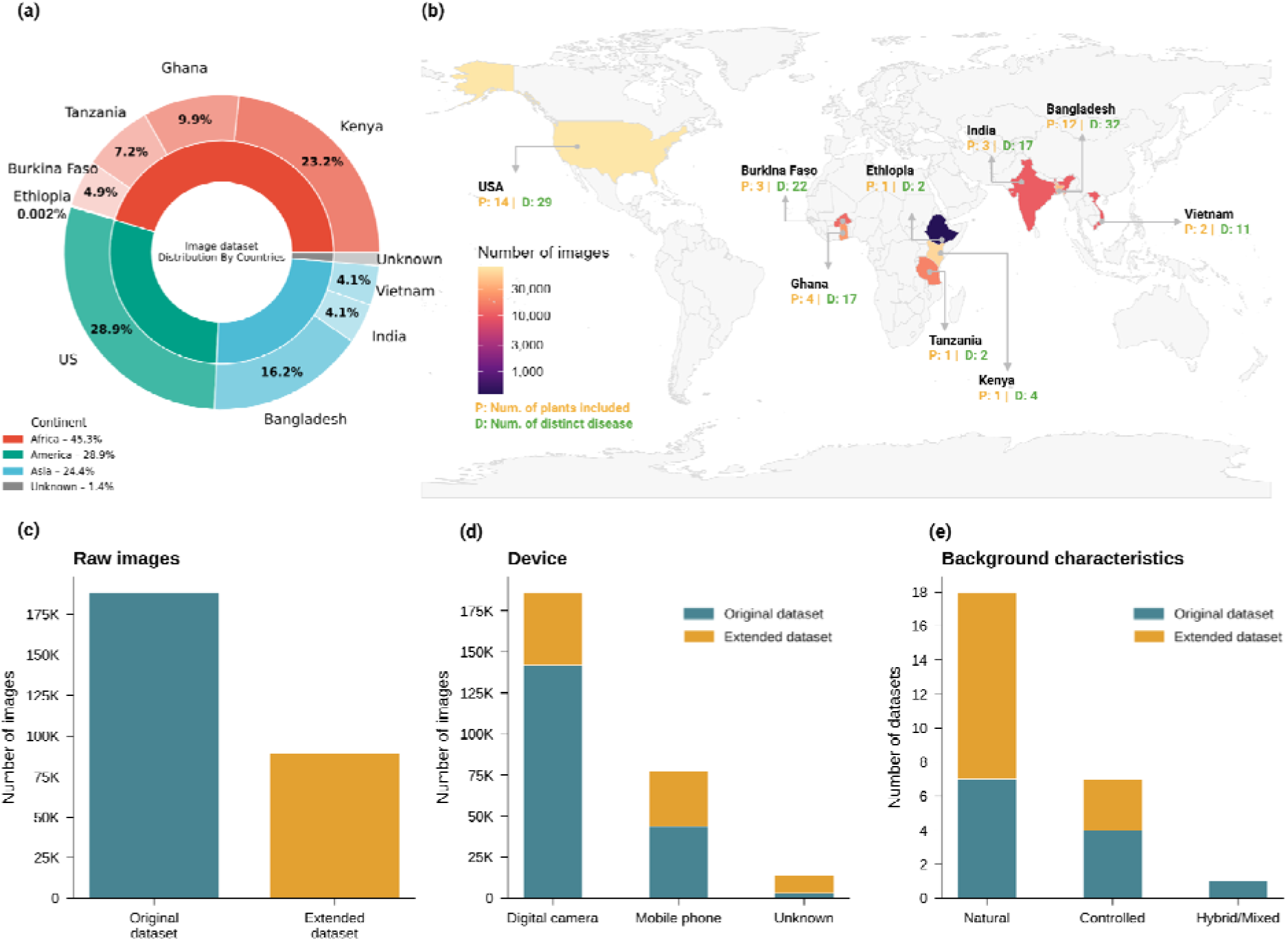
Geographic coverage and dataset composition of LeafNet2.0. **(a)** Distribution of images across continents (inner ring) and countries (outer ring), illustrating the relative contribution of each region to the dataset. **(b)** Global distribution of crop species and plant diseases represented in LeafNet2.0 across different geographic regions. Labels indicate the number of plant species (P) and distinct disease categories (D) represented in each country. **(c–e)** Dataset composition of LeafNet2.0, including comparison of the number of raw images between the original and extended datasets, distribution of image acquisition devices, and distribution of background environments represented in the dataset, including natural, controlled, and hybrid/mixed settings, where contributing datasets contain multiple background environment types.

To create a more standardized dataset, a quality control process was conducted using several criteria. Samples with unclear or barely visible disease symptoms were excluded because they provide limited diagnostic information. Most excluded samples contained disease symptoms that occupied only a negligible portion of the leaf area, making them difficult to discern through visual inspection. Consequently, the characteristic symptom features described in the corresponding reference literature could not be confidently verified. Images not centered on leaf tissue, including datasets containing fruits, stems, or multiple plant organs alongside leaves, were also excluded to maintain a consistent leaf focus across samples

Leaves exhibiting multiple diseases were furthermore excluded. For example, a tomato leaf may exhibit the concentric necrotic lesions characteristic of early blight while simultaneously displaying white powdery growth typical of powdery mildew. Such symptom combinations are biologically plausible under field conditions, as primary pathogen infection can alter host physiology or suppress defense responses, thereby increasing susceptibility to subsequent infections (Abdullah, 2017). However, the existing source materials generally reported only a single confirmed diagnosis and rarely documented whether additional symptoms represented true co-infection, secondary colonization, or independent disease processes. Assigning multiple disease labels in these cases would therefore require assumptions beyond the available evidence and could introduce annotation uncertainty. To preserve label reliability and ensure that each annotation remained traceable to a documented diagnosis, only images with a clearly supported disease label were retained.. Incomplete or corrupted files were additionally discarded. All retained images were resized to 224 × 224 pixels to ensure compatibility with widely used pre-trained computer vision models while preserving diagnostically relevant leaf features.

Following the above process, 52,136 images were manually removed, resulting in a final collection of 255,825 leaf disease images in LeafNet2.0. To the best of our knowledge, this currently represents the largest publicly available large-scale dataset dedicated specifically to leaf disease recognition.

### B. Caption Construction

The annotation process was conducted across 197 crop–disease classes. Trained annotators were assigned subsets of crop–disease pairs and performed two independent tasks: (1) standardizing metadata annotations in the extended dataset to ensure consistency with the original dataset structure, and (2) constructing class-level captions using literature-grounded references in conjunction with the standardized metadata fields.

For metadata annotation, each image was annotated with pathogen type, scientific name, leaf shape, disease status, and background condition. Pathogen names and scientific classifications were primarily obtained from the original publications or source datasets from which the extended images were collected. When source datasets did not provide sufficient metadata, annotations were verified using established plant pathology references and published literature. Leaf shape annotations were visually inspected and compared against published references to maintain morphological consistency across crop species. Disease status and background conditions were manually annotated through visual assessment of each image to capture both symptom presentation and environmental context. Because many classes were aggregated from a single source, image backgrounds and acquisition conditions were often visually consistent within a class. However, when a class contained heterogeneous backgrounds or multiple imaging conditions, these variations were documented at the image level to preserve contextual fidelity during caption construction.

To reduce biologically inaccurate or overly generic outputs from large language models (LLMs), human validation was integrated at the earliest stage of the captioning pipeline rather than introduced as a post hoc correction step. Annotators first constructed concise, literature-grounded reference descriptions to preserve diagnostically relevant agricultural terminology and symptom characteristics (Fig. 1b), which were subsequently used to guide generation of the final captions. To achieve this, annotators systematically reviewed peer-reviewed literature retrieved through Google Scholar using structured queries combining crop name, disease name, and symptom-related terms. To maintain biological reliability and terminological consistency, references were restricted to established scientific publishers, including Springer Nature, Elsevier (ScienceDirect), and other comparable peer-reviewed sources. Agricultural extension reports were additionally consulted as supplementary references because they often provide field-oriented descriptions of symptom progression, phenotypic variability under natural growing conditions, and visually observable characteristics that are frequently underrepresented in formal pathology literature. These resources, however, were incorporated only when their descriptions were consistent with and traceable to primary academic references.

Annotators then answered a standardized set of symptom-oriented questions targeting diagnostically relevant visual traits, including lesion morphology, color transition patterns, spatial distribution across the leaf surface, and structural deformation of infected tissues, with the clear guiding questions, shown in Table 2. Each symptom category was described for two representative stages of disease progression: early and late. Intermediate stages were intentionally excluded because they are frequently ambiguously defined, visually overlapping, and inconsistently represented in leaf-level imagery across datasets and studies. In contrast, early and late stages correspond to clinically and visually distinguishable phases of infection that are more consistently documented in plant pathology literature. Annotators therefore synthesized information from the reviewed sources into concise one- to two-sentence descriptions for each crop–disease pair and disease stage, ensuring that the resulting captions remained both biologically grounded and semantically standardized across the dataset.

**Table 1.**
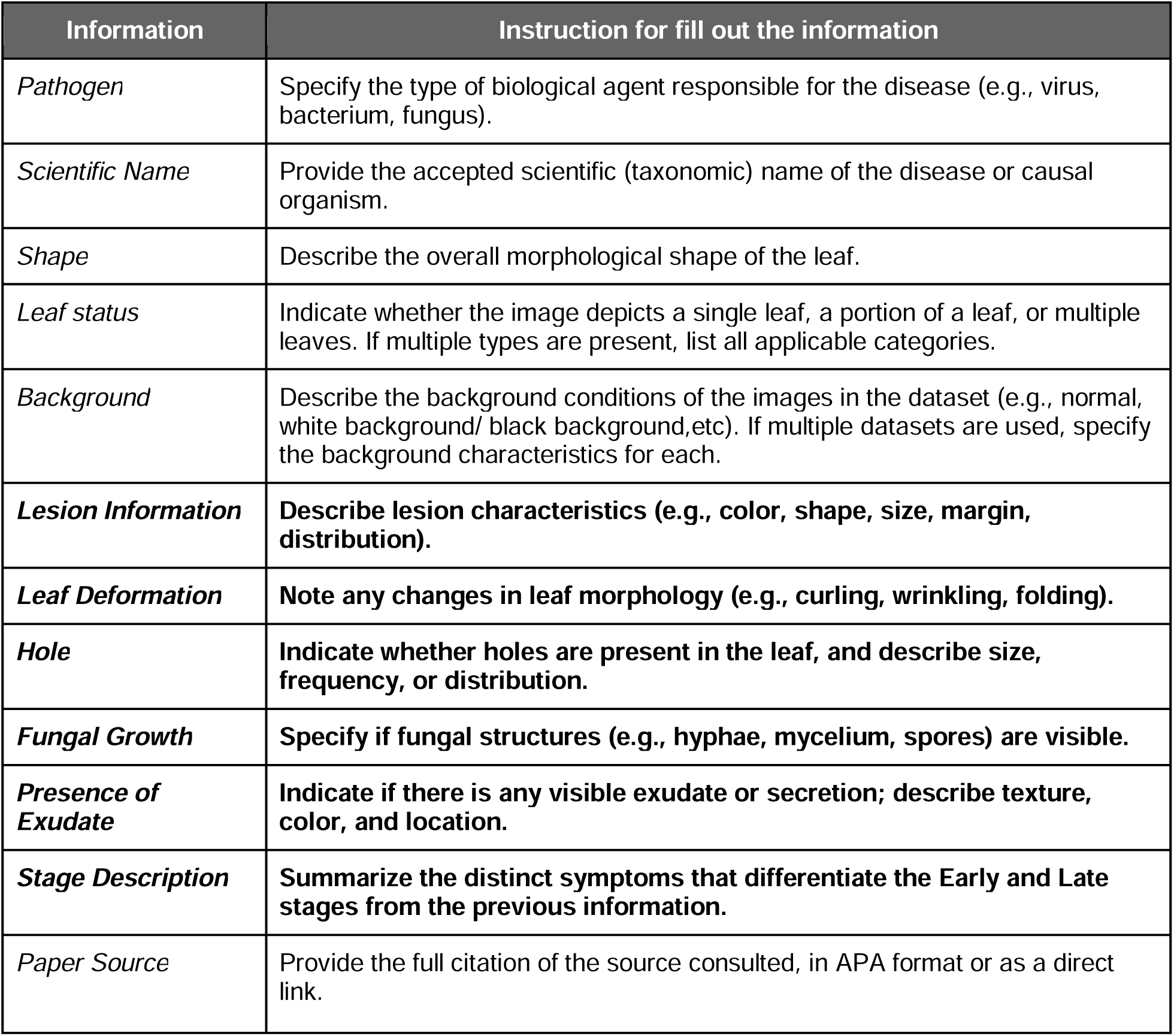
Redefined annotation template used to standardize the recording of diagnostically relevant leaf disease symptoms across crop–disease classes. Fields shown in bold correspond to attributes directly derived from literature-grounded symptom descriptions.

**Table 2.** Criteria and instruction questions applied by expert reviewers during manual dataset validation. Samples for which any criterion received a “No” response were flagged and subjected to additional review or revision.

| Criteria | Instruction question |
| --- | --- |
| Crop | Does the visible leaf morphology match the labeled crop species? |
| Disease | Are the visible symptoms consistent with the labeled disease? |
| Primary Symptom | Does the caption describe disease-specific symptom characteristics, rather than generic visual cues? |
| Disease Stage | If a disease stage is specified, is the assigned stage visually supported by symptom severity, lesion size, and spatial spread? |
| Revision criteria | **If any of the above criteria are not satisfied, the sample should be flagged for revision or exclusion, except in cases where disease stage information is intentionally absent.** |

Before the second stage of caption generation, as the human-authored may have subjective error, all human-authored reference captions underwent parallel quality control using LLMs to ensure that the caption is clear, relevant and descriptive enough. Here we use 3 state-of-the-art models Google DeepMind Gemini Pro 2.5 (Comanici et al., 2025), OpenAI GPT-4 (OpenAI et al., 2023), and Anthropic Claude 3.5 (Anthropic, n.d.), reduced dependence on model-specific linguistic and interpretative biases in single-model judgment. Each caption was independently evaluated for clarity, relevance to visible disease symptoms, and descriptive completeness using a standardized prompting framework (Fig. 3) and a five-point scoring scale.

**Fig. 3.**
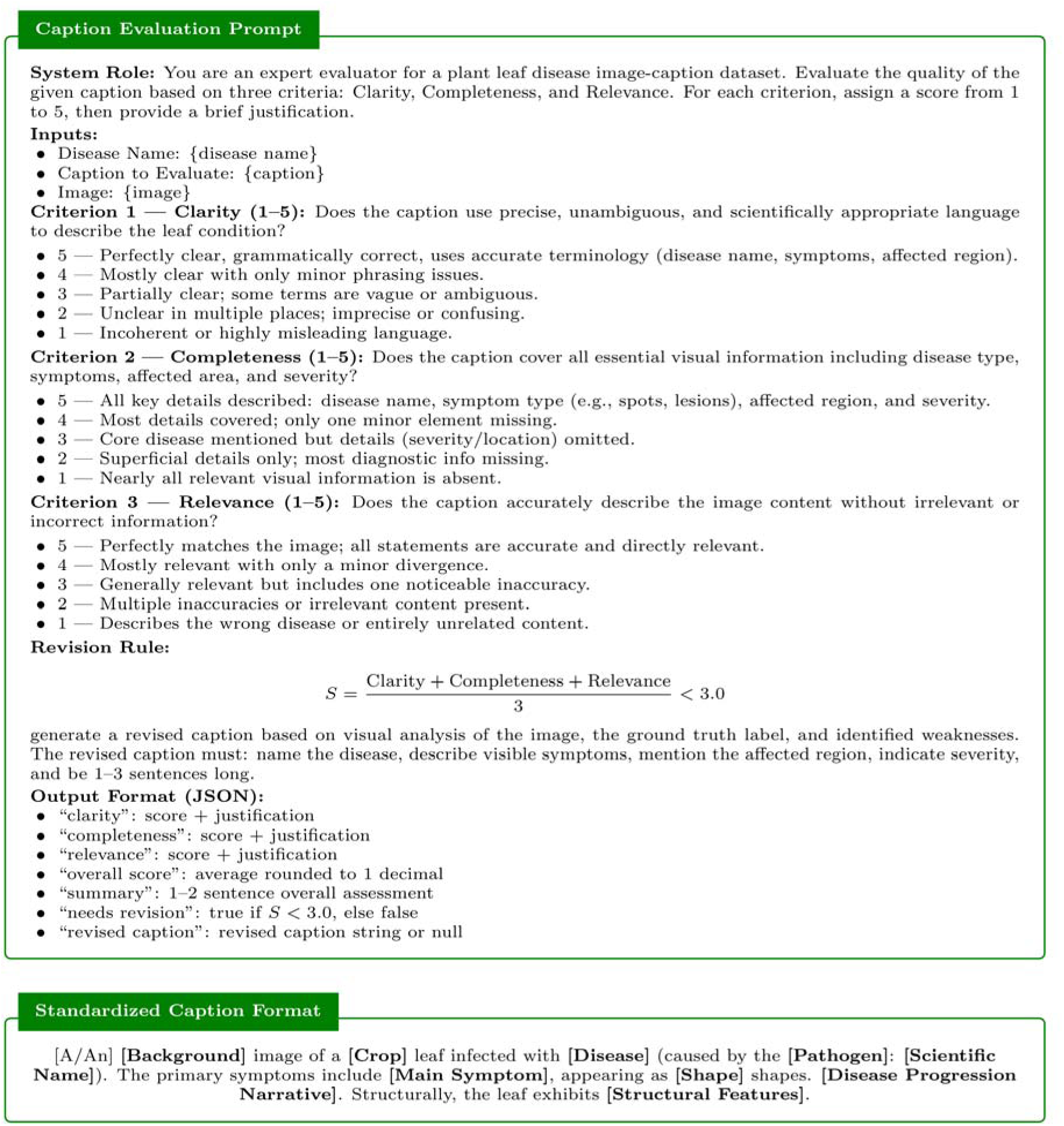
LLM-guided caption evaluation and standardized caption construction framework used in LeafNet2.0. The upper panel shows the evaluation prompt used to assess generated captions based on clarity, completeness, and relevance, together with the corresponding revision criteria and structured output format. The lower panel presents the standardized caption template designed to maintain biological consistency and terminological uniformity across crop–disease classes.

Captions receiving a score of three or lower from at least one of the three models were flagged for revision. For these cases, alternative captions were regenerated using the same models under the original evaluation criteria. The revised outputs were then examined by domain experts, who selected the caption that most accurately reflected the visible symptoms and disease characteristics. This process resulted in a set of consistent and biologically grounded reference captions for the subsequent annotation stage.

Using these expert-validated reference captions as class-level baselines, the second stage focused on generating standardized image-specific captions across the full dataset. All images were processed using OpenAI GPT-4, which was selected following preliminary comparisons with Gemini Pro 2.5 and Claude 3.5. GPT-4 produced more consistent outputs in preserving disease-specific terminology, aligning visible symptoms with the reference descriptions, and maintaining stable caption structures across heterogeneous imaging conditions. This observation is consistent with previous studies reporting strong instruction-following and structured text generation capabilities of GPT-4 in scientific and multimodal applications (OpenAI et al., 2023; Bubeck et al., 2023). Because this stage focused primarily on standardizing image-level descriptions from expert-defined references rather than generating new biological interpretations, GPT-4 was used to improve consistency and formatting uniformity across the dataset.

In the second stage, 255,855 synthetic captions were generated by processing all images using GPT-4, with the corresponding reference captions provided as contextual guidance. The model produced expanded captions integrating image context with descriptions of visible disease symptoms, informed by both the reference captions and the visual characteristics of each image. All generated captions followed a standardized structure (Fig. 3), describing image context, primary symptoms, and, when visually distinguishable, disease stage and progression. Disease stage information was included only when supported by visible evidence, as many images did not exhibit clear stage-specific characteristics.

### C. Validation

#### Expert Validation

To address substantial class imbalance across crop–disease categories, a stratified sampling strategy was adopted during manual quality assessment. Rare classes were fully reviewed to preserve low-frequency symptom variability and minimize the risk of overlooking annotation errors in underrepresented categories, whereas larger classes were evaluated through capped random subsampling to maintain feasible large-scale inspection while preserving representative intra-class diversity. Specifically, classes containing fewer than 50 images were fully reviewed; classes with 50–299 images were sampled with 25 images; classes with 300–1,999 images were sampled with 35 images; and classes containing 2,000 or more images were sampled with 50 images. Overall, 6,361 image–caption pairs were included in the manual quality assessment and benchmark evaluation.

Each sampled image was systematically examined to verify consistency between the visual content and its associated metadata, including background context, crop species, disease label, primary symptom, and disease stage. Detailed verification criteria are summarized in Table 3. Expert evaluators assigned binary scores to each criterion, where 1 indicated that the annotation satisfied the verification standard and 0 indicated non-compliance. Disease stage annotations were treated separately because symptom progression was not always visually distinguishable across all images. Samples receiving a score of 0 in any mandatory criterion were flagged as unqualified, and the corresponding class underwent further review and revision to improve annotation consistency and biological reliability.

**Table 3.** Quantitative benchmark results for all evaluated models on LeafBench 2.0, including 7 CLIP-based models, 7 generative vision–language models, and 2 proprietary models. Orange values indicate the best-performing model for each task.

| Model | DI | PC | CSI | SI | HDC | SNC | LI | LSD | DSC | Avg |
| --- | --- | --- | --- | --- | --- | --- | --- | --- | --- | --- |
| *Dual-Encoder Models* |  |  |  |  |  |  |  |  |  |  |
| SigLIP2 | 20.71 | 5.65 | 26.61 | 27.32 | 12.43 | 20.65 | 26.20 | 7.72 | 53.76 | 22.34 |
| SigLIP | 24.07 | 18.30 | 60.69 | 26.97 | 92.04 | 25.98 | 33.64 | 61.27 | 52.72 | 43.96 |
| BioCLIP | 41.12 | 18.90 | 71.00 | 32.94 | 20.69 | 31.00 | 26.99 | 26.04 | 45.78 | 34.94 |
| CLIP | 27.50 | 13.29 | 62.37 | 25.01 | 77.86 | 28.07 | 33.23 | 40.31 | 52.69 | 40.04 |
| OpenAI-CLIP | 28.85 | 19.68 | 59.96 | 26.44 | 87.27 | 26.32 | 37.44 | 44.46 | 50.84 | 42.36 |
| SCOLD | 62.90 | 19.49 | 68.18 | 59.46 | 39.63 | 27.26 | 40.50 | 41.77 | 47.76 | 45.22 |
| AgriCLIP | 65.10 | 22.40 | 71.20 | 62.50 | 45.60 | 29.80 | 43.00 | 48.50 | 51.40 | 48.86 |
| *Open Vision Language Models* |  |  |  |  |  |  |  |  |  |  |
| LFM2 1.6B | 29.81 | 43.80 | 58.50 | 29.31 | 74.83 | 23.11 | 27.78 | 65.79 | 61.00 | 45.99 |
| Qwen2.5 3B | 34.43 | 44.81 | 62.01 | 35.84 | 83.52 | 29.90 | 34.98 | 74.39 | 48.55 | 59.00 |
| SmolVLM2 | 36.43 | 36.43 | 44.80 | 34.51 | 42.40 | 23.44 | 27.63 | 63.50 | 73.50 | 42.52 |
| moondream2 | 26.40 | 51.00 | 71.76 | 27.45 | 71.47 | 21.46 | 29.83 | 34.02 | 68.97 | 44.71 |
| Qwen3-VL-4B | 50.86 | 55.76 | 76.21 | 54.79 | 74.98 | 41.91 | 44.40 | 85.16 | 56.13 | 60.02 |
| Gemma4-5B | 39.07 | 46.19 | 37.52 | 36.85 | 57.12 | 24.04 | 33.30 | 88.41 | 47.99 | 45.61 |
| Gemma4-8B | 47.57 | 42.63 | 48.06 | 43.90 | 70.01 | 30.69 | 34.96 | 92.76 | 55.09 | 51.84 |
| *Proprietary Vision Language Models* |  |  |  |  |  |  |  |  |  |  |
| GPT-4o | 64.30 | 58.20 | 74.10 | 52.40 | 93.50 | 40.60 | 48.90 | 88.20 | 62.30 | 64.72 |
| Gemini 2.5 Pro | 67.80 | 60.50 | 76.80 | 55.90 | 95.70 | 44.20 | 52.10 | 91.40 | 65.60 | 67.78 |

**Abbreviations:** DI = Disease Identification; PC = Pathogen Classification; CSI = Crop Species Identification; SI = Symptom Identification ; HDC = Healthy-Diseased Classification ; SNC = Scientific Name Classification; LI = Lesion Identification; LSD = Leaf Symptom Detection; DSC = Disease Severity Classification

#### Benchmarking Model

To enable systematic evaluation of fine-grained plant disease understanding in vision–language models, we developed LeafBench 2.0, a multiple-choice visual question answering benchmark derived from the LeafNet 2.0 dataset. Each benchmark sample consists of a leaf image, a task-specific question, and four candidate answers containing one correct choice and three distractors. The benchmark covers nine complementary tasks related to plant disease analysis: disease identification (DI), pathogen classification (PC), crop species identification (CSI), symptom identification (SI), healthy–diseased classification (HDC), scientific name classification (SNC), lesion identification (LI), leaf symptom detection (LSD), and disease severity classification (DSC). Together, these tasks evaluate the ability of models to identify disease categories, infer causal agents, identify crop species, and interpret visual disease symptoms from leaf images (Fig. 1c). Detailed task definitions and representative examples are provided in Tables S3–S4.

Unlike conventional image classification benchmarks that primarily assess category-level recognition, LeafBench 2.0 evaluates the ability of models to discriminate among visually similar diseases, subtle symptom variations, and closely related pathogen manifestations under heterogeneous field conditions. Because the benchmark was developed directly from the LeafNet 2.0 dataset, it is also expected to preserve substantial real-world variability in imaging devices, environmental backgrounds, illumination conditions, symptom progression, and disease severity, thereby providing a more realistic evaluation setting for agricultural vision systems.

We evaluated benchmark performance using 16 vision–language models spanning multiple architectural paradigms, including 7 CLIP-based models: SigLIP2 (Tschannen et al., 2025), SigLIP (Zhai et al., 2023), BioCLIP (Stevens et al., 2023), CLIP (Radford et al., 2021), OpenAI-CLIP (OpenAI, n.d.), SCOLD (Nguyen Quoc et al., 2025), and AgriCLIP (Gao et al., 2025); 7 generative vision–language models: LFM2 1.6B (Amini et al., 2025), Qwen2.5 3B (Hui et al., 2024), SmolVLM2 (Marafioti et al., 2025), moondream2 (Vikhyatk, n.d.), Qwen3-VL-4B (Bai et al., 2025), Gemma4-5B (Gemma Team, 2025), and Gemma4-8B (Gemma Team, 2025); and 2 proprietary models, GPT-4o (OpenAI et al., 2023) and Gemini 2.5 Pro (Comanici et al., 2025). Model performance was assessed using classification accuracy across all benchmark tasks, and the corresponding results are presented in Table 3.

## Data Records

LeafMD consists of two complementary datasets: LeafNet 2.0 and LeafBench 2.0.

The LeafNet 2.0 dataset is organized into three main components: an **images/** directory containing all leaf images, an **image_caption_pair/** directory containing tabular image–caption annotations, and a metadata/ directory containing auxiliary information used during dataset construction and validation (Fig. 4a).

**Fig. 4.**
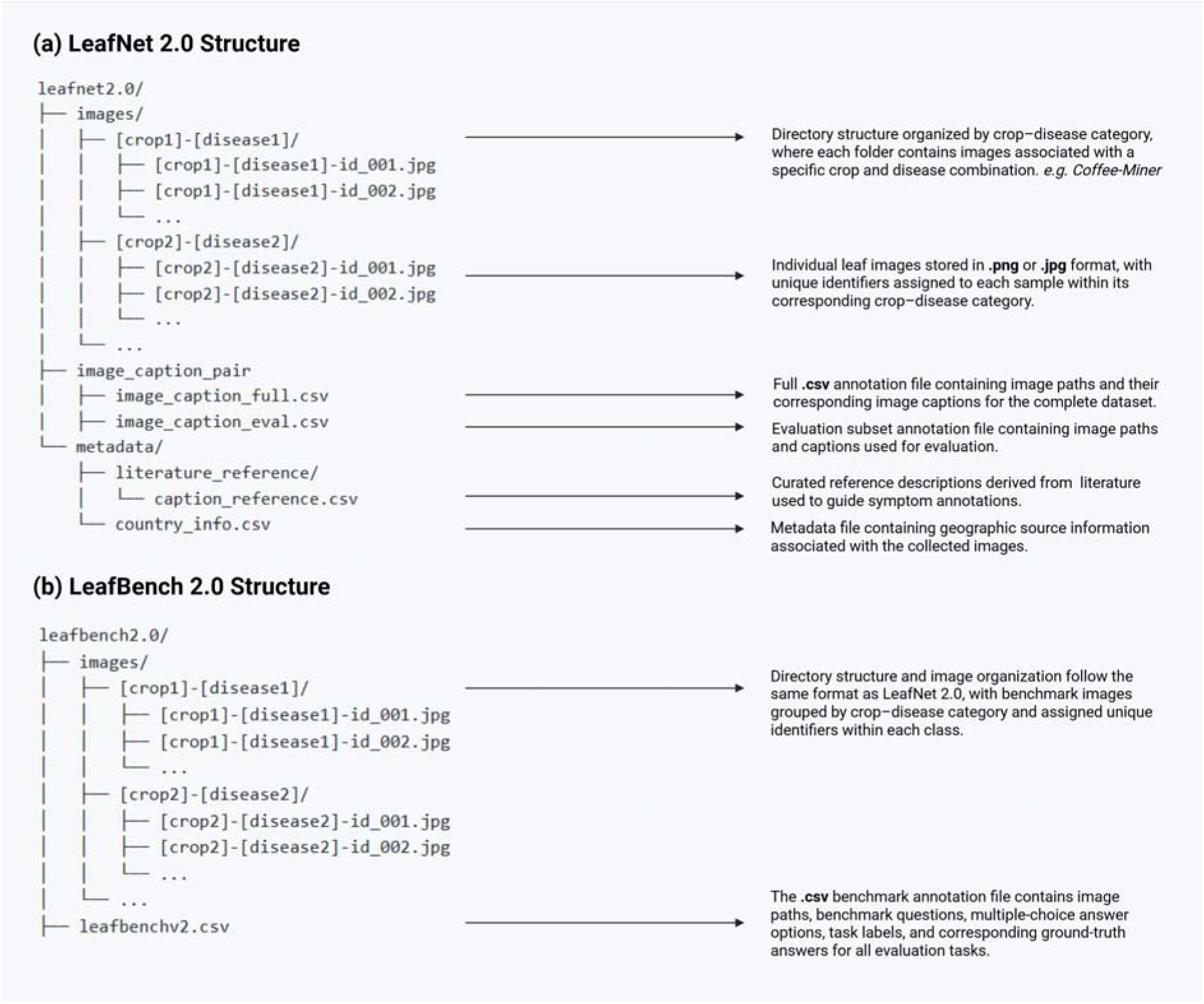
Directory structure and annotation organization of LeafNet 2.0 and LeafBench 2.0. (a) Organization of the LeafNet 2.0 dataset, including crop–disease-specific image folders, image–caption annotation files, and metadata files containing literature-based symptom references and geographic source information. (b) Organization of the LeafBench 2.0 benchmark, following the same crop–disease directory structure as LeafNet 2.0, together with the benchmark annotation file containing image paths, benchmark questions, multiple-choice answer options, task labels, and corresponding ground-truth answers for all evaluation tasks.

The images/ directory contains crop–disease-specific subdirectories named using the format *[crop]-[disease]*, with images stored in **.jpg** or **.png** format and indexed using the standardized naming convention *[crop]-[disease]-id_[ID]*.

The **image_caption_pair/** directory contains two annotation files: **image_caption_full.csv**, which includes the complete set of **image–caption pairs**, and **image_caption_eval.csv**, which contains the evaluation subset used for manual validation and benchmark assessment.

The **metadata/** directory includes **caption_reference.csv**, containing literature-based symptom references used for caption generation, and **country_info.csv**, which provides geographic source information associated with the collected images.

Similarly, the LeafBench 2.0 benchmark consists of an **images/** directory and a benchmark annotation file, leafbenchv2.csv, as illustrated in Fig. 4b. The **images/** directory follows the same hierarchical organization as LeafNet 2.0, where images are grouped into crop–disease-specific subdirectories named using the format *[crop]-[disease]*. Image files are stored in .jpg or .png format and named using the standardized convention *[crop]-[disease]-id_[ID]*.

The **leafbenchv2.csv** file contains the complete benchmark annotations used for model evaluation. Each row corresponds to a single benchmark instance and includes the image path, benchmark task category, question, four multiple-choice answer options, and the corresponding ground-truth answer. The benchmark covers all nine evaluation tasks and is directly derived from LeafNet 2.0 image–caption pairs, preserving the same variability in symptom appearance, imaging conditions, and field backgrounds observed in real-world agricultural environments.

## Technical Validations

The results of the two-step validation process are summarized in Fig. 5.

**Fig. 5.**
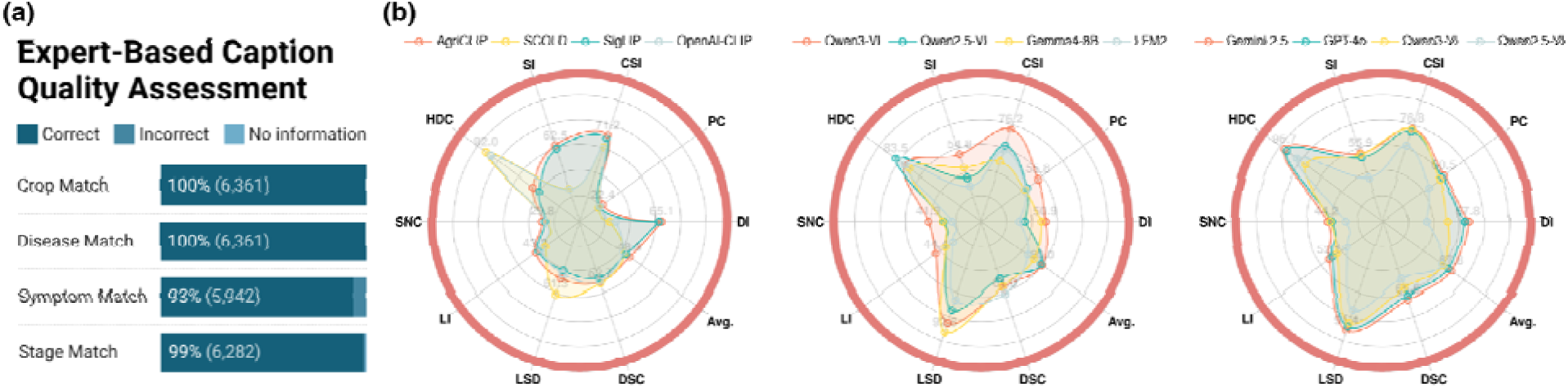
Benchmark evaluation on LeafBench 2.0. (a) Expert-based quality assessment of image–caption pairs used for benchmark construction.(b) Radar plots showing task-wise performance of representative CLIP-based, generative, and proprietary vision–language models across the nine benchmark tasks.

### Expert Evaluation

Fig.5a summarizes the results of the two-step expert validation described in Table 3. Crop species and disease labels achieved 100% agreement with the ground-truth annotations across all 6,361 evaluated image–caption pairs, indicating that the caption generation pipeline consistently preserved the primary biological identity of each sample. This high agreement is particularly important because crop identity and disease class constitute the core semantic information required for downstream plant disease recognition and vision–language learning tasks.

The disease stage annotations also achieved high agreement during manual validation. Among the 6,361 evaluated image–caption pairs, 6,319 images contained sufficient visual information to determine disease stage, whereas the remaining samples were labeled as “No information” because the images did not exhibit clearly distinguishable stage-specific characteristics. Of the images with identifiable stage information, 99% were consistent with expert annotations, while fewer than 1% showed stage mismatches. These results indicate that disease progression information can be reliably incorporated into large-scale image–caption annotations when supported by visible symptom evidence.

Among the evaluated samples, 7% required revision during manual inspection because the generated symptom descriptions were not fully consistent with the visible symptoms in the corresponding images. The most common issues involved captions that remained overly generic and therefore insufficiently representative of the disease phenotype, or symptom descriptions that only partially matched the visible disease characteristics. These inconsistencies primarily reflected the intrinsic visual ambiguity of field-acquired plant disease imagery rather than systematic annotation failures. In many cases, symptoms appeared at early or intermediate stages, were partially occluded, overlapped with environmental stress responses, or occupied only a small region of the image, making fine-grained symptom interpretation challenging even for expert annotators. However, these inconsistencies affected only a small proportion of the dataset, while crop species and disease labels remained fully consistent with the ground-truth annotations across all evaluated samples, indicating that the overall caption generation and validation pipeline remained highly reliable for large-scale vision–language modeling applications.

### Benchmark Evaluation

As shown in Fig. 5b, there is a clear difference in task specialization across model families. Generally, proprietary vision–language models exhibited the most balanced performance profiles across benchmark tasks, suggesting stronger generalization across diverse disease phenotypes, imaging conditions, and symptom complexities. In contrast, dual-encoders modelsand open vision–language models showed greater task-specific variability. Agriculture-adapted dual-encoder models, particularly AgriCLIP and SCOLD, demonstrated comparatively strong performance in crop species identification (CSI) and symptom-related tasks, indicating that domain-specific pretraining substantially improves fine-grained agricultural representation learning. Open vision–language models generally showed more balanced performance in healthy–diseased classification (HDC) and leaf symptom detection (LSD), suggesting stronger integration of global contextual information and symptomatic reasoning compared with contrastive dual-encoders architectures.

Table 3 further confirms the superior performance of proprietary models, with Gemini 2.5 Pro achieving the highest overall accuracy (67.78%), followed by GPT-4o (64.72%). Task-wise performance revealed substantial variability in benchmark difficulty. CSI and HDC were generally the most tractable tasks, with several models achieving accuracies above 90%. Gemini 2.5 Pro achieved the highest CSI performance (76.80%), whereas GPT-4o obtained the strongest HDC performance (93.50%). In contrast, pathogen classification (PC), scientific name classification (SNC), and lesion identification (LI) remained considerably more challenging across architectures, likely because these tasks require discrimination between visually similar diseases and subtle lesion characteristics that differ only in localized texture, boundary structure, or color variation. Among open-source approaches, AgriCLIP achieved the strongest performance within the dual-encoder models category, particularly for disease identification (65.10%) and symptom identification (62.50%), outperforming several larger general-domain models. Qwen3-VL-4B achieved the strongest overall performance among open-source generative models (60.02% average accuracy). Interestingly, LSD achieved relatively high performance across multiple architectures, with Gemma4-8B reaching the highest score (92.76%), indicating that identifying symptomatic regions is generally easier than assigning precise pathogen identities or scientific disease categories.

Overall, the results suggest that LeafBench 2.0 captures meaningful fine-grained complexity in real-world plant disease interpretation rather than only coarse disease identification. While most models performed well on broader tasks such as crop species identification and healthy–diseased classification, performance dropped noticeably for tasks requiring subtle pathological reasoning, including pathogen classification, lesion identification, and scientific name classification. The clear performance differences across architectures, particularly between general-domain and agriculture-adapted models, also indicate that the benchmark is sensitive to domain-specific agricultural knowledge. Together, these findings suggest that LeafBench 2.0 provides a realistic and sufficiently challenging evaluation setting for future vision–language models in plant disease analysis.

## Usage Note

LeafNet 2.0 is particularly suitable for studying multimodal disease representations under heterogeneous agricultural conditions, as the dataset spans multiple crop systems, geographic regions, environmental backgrounds, imaging conditions, and disease stages. Because symptom expression often varies across climates, cultivation practices, and developmental stages, the dataset may be useful for evaluating model robustness under substantial visual and biological distributional shifts. The integration of disease stage annotations further enables analyses of symptom progression and temporal changes in disease manifestation, which remain underexplored in existing plant pathology datasets.

Furthermore, the image–caption structure of LeafNet 2.0 supports downstream tasks that require both visual recognition and symptom-level interpretation. Potential applications include symptom-aware image retrieval, multimodal embedding learning, disease-aware caption generation, visual question answering, and pretraining of agricultural foundation models. For example, models trained on LeafNet 2.0 may learn associations between visual symptoms such as chlorotic halos, necrotic lesions, vein discoloration, or powdery fungal growth and their corresponding pathological descriptions, enabling biologically informed disease representations beyond categorical classification alone.

To systematically evaluate these capabilities, LeafBench 2.0 provides a complementary evaluation framework for assessing fine-grained disease understanding in vision–language models. In contrast to conventional benchmarks focused primarily on disease classification accuracy, LeafBench 2.0 evaluates multiple levels of pathological reasoning, including lesion characterization, symptom interpretation, pathogen attribution, disease stage recognition, and disease severity assessment. These tasks may be particularly useful for examining whether multimodal models capture biologically meaningful symptom features rather than relying predominantly on superficial visual correlations.

Together, LeafNet 2.0 and LeafBench 2.0 are expected to foster the development of more interpretable and transferable AI systems for plant pathology. Potential applications include extension-oriented diagnostic support systems, symptom-based retrieval of biologically similar disease cases across crops and geographic regions, automated pathology description generation from field-acquired imagery, and multimodal foundation models capable of adapting across heterogeneous agricultural environments.

Several characteristics of the dataset should nevertheless be considered during downstream use. First, although the image–caption pairs were generated through an expert-guided pipeline and manually validated, symptom interpretation in field-acquired imagery remains inherently ambiguous in some cases. A small proportion of samples contained partially inconsistent symptom descriptions, particularly when symptoms were subtle, spatially localized, partially occluded, or visually confounded with environmental stress responses. Importantly, these inconsistencies were concentrated primarily at the symptom-description level rather than crop or disease identity, which remained highly consistent throughout validation.

Second, disease stage annotations were included only when stage-specific characteristics were visually distinguishable. Consequently, progression labels should not be interpreted as exhaustive representations of temporal disease development. Third, because the dataset was aggregated from heterogeneous public and partner sources, substantial variation remains in image quality, illumination conditions, camera devices, and background complexity across categories. While this variability increases realism for field deployment scenarios, it may also introduce additional challenges for highly controlled benchmarking settings.

Finally, the dataset construction process involved extensive filtering and manual quality control to remove duplicated, corrupted, low-quality, or biologically inconsistent samples. As a result, approximately 53,000 candidate images were excluded during curation. Although this process improved overall annotation reliability and dataset consistency, it also reflects a broader limitation of large-scale plant disease data collection, namely the substantial loss of potentially usable field imagery due to inconsistent annotations, poor visual quality, or insufficient pathological information.

The benchmark results further suggest that current vision–language models still rely heavily on coarse visual cues and remain limited in fine-grained pathological reasoning. Most evaluated architectures performed substantially better on crop species identification and healthy–diseased classification than on pathogen classification, scientific name classification, or lesion identification, where visually similar diseases often differ only in subtle lesion morphology or localized symptom patterns. Interestingly, agriculture-adapted models such as AgriCLIP and SCOLD consistently outperformed several larger general-domain architectures on symptom-related tasks, indicating that domain-specific agricultural pretraining contributes meaningful improvements in disease-aware representation learning. These findings suggest that LeafBench 2.0 may serve not only as a benchmarking resource, but also as a useful platform for studying multimodal reasoning, agricultural domain adaptation, and biologically informed representation learning in future foundation models.

Future extensions of LeafNet 2.0 may further improve both the biological coverage and multimodal richness of the dataset. One important direction is the inclusion of additional disease progression stages, longitudinal observations, and temporal sequences that enable more comprehensive modeling of symptom evolution through time. Expanding geographic coverage to underrepresented agricultural regions, particularly in low-resource tropical systems, may also improve the study of distributional shifts and cross-regional generalization in plant disease models. The dataset can be expanded through an open, online community data-sharing and contribution portal designed to facilitate data contributions from researchers, collaborators, farmers, and other stakeholders, thereby improving both dataset scale and representation.

At the same time, future work may focus on reducing the substantial proportion of field-acquired imagery discarded during dataset curation. Developing weakly supervised annotation strategies, uncertainty-aware labeling methods, automated quality assessment systems, and human–AI collaborative validation pipelines may therefore help preserve biologically relevant but imperfect samples while maintaining annotation quality at scale. Beyond image–text pairs, future versions of the dataset may additionally incorporate complementary modalities such as canopy-level observations, environmental measurements, management information, genomic data, or laboratory diagnostic records. Integrating these heterogeneous data sources could support the development of more biologically informed agricultural foundation models capable of linking visual symptoms with environmental drivers and pathogen dynamics across scales.

In parallel, future extensions of LeafBench 2.0 may move toward more complex reasoning tasks involving causal interpretation, uncertainty estimation, differential diagnosis, and multi-disease recognition under realistic field conditions. Future evaluation settings could also examine how models perform on unseen crops, emerging pathogens, or disease distributions shifted across geographic regions, where symptom expression and field conditions may differ substantially from the training data. Together, these efforts may help move plant pathology AI beyond highly controlled benchmark settings toward models that can operate more reliably across the variability, uncertainty, and complexity of real agricultural environments.

## Data availability

LeafMD are publicly available and can be accessed from the following link: https://huggingface.co/collections/enalis/leafmd

## Code availability

All details of the code, packages, and implementation are available at https://github.com/EnalisUs/LeafBench. All experiments were conducted in Python 3.10 with torch v2.7.0, openCV v4.11.0.86, and transformers v4.51.3. Additional essential packages include accelerate (1.8.1), torchvision (0.22.0), and peft (0.15.0). All figures and radar plots were generated using Microsoft PowerPoint, BioRender and seaborn v0.13.0. Standard vision backbones, including ViT-B/16, VGG16, MobileNetV3-Small, DenseNet121, EfficientNet-B0, and EfficientNet-V2S, were implemented using the timm (PyTorch Image Models) library (https://github.com/huggingface/pytorch-image-models). Furthermore, agricultural-specific and generative VLMs were obtained from their respective repositories: AgriCLIP, BioCLIP (https://github.com/microsoft/BioCLIP), SCOLD (https://huggingface.co/enalis/scold), MoonDream2 (https://github.com/vikhyat/moondream), and the Qwen (https://github.com/QwenLM/Qwen-VL) and Gemma4 (https://github.com/google/gemma-vlm) model families.

## Author Contributions

**Trang V. Nguyen:** Data curation, Formal analysis, Validation, Visualization, Writing – original draft. **Khang Nguyen Quoc:** Conceptualization, Data curation, Data modeling, Validation, Visualization, Writing – review & editing. **David Harwath:** Supervision, Writing – review & editing. **Luyl-Da Quach:** Data curation & Supervision **Phuong D. Dao:** Conceptualization, Project administration, Supervision, Writing – review & editing, Funding acquisition.

## Competing Interests

The authors declare no competing interests.

## Funding

This work was supported by Dr. Phuong D. Dao’s startup fund provided by The University of Texas at Austin. Khang Nguyen Quoc was supported by the Hyundai Motor Chung Mong-Koo Foundation Global Scholarship (GSS-25-02120).

## Notes

https://huggingface.co/collections/enalis/leafmd

### Competing Interest Statement

The authors have declared no competing interest.

### Summary of Updates

Author name updated due to typo

