## Supplementary Information for "A multiregional image–text dataset and benchmark for vision-language modeling of plant diseases"

**SUPPLEMENT DOCUMENTS**

**Table S1 | Summary statistics of crops and diseases included in the dataset, with sample counts**

| **Crop** | **Disease** | **Sample Number** |
| --- | --- | --- |
| Apple | Scab | 5543 |
| Apple | Black rot | 621 |
| Apple | Cedar apple rust | 364 |
| Apple | Healthy | 6294 |
| Apple | Powdery mildew | 1271 |
| Apple | Frog eye leaf spot | 3346 |
| Apple | Rust | 1957 |
| Cashew | Anthracnose | 1672 |
| Cashew | Healthy | 1045 |
| Cashew | Leaf miner | 1378 |
| Cashew | Rust | 1682 |
| Cassava | Bacterial blight | 2614 |
| Cassava | Brown spot | 1481 |
| Cassava | Green mite | 1015 |
| Cassava | Healthy | 1193 |
| Cassava | Mosaic | 1205 |
| Cherry | Healthy | 854 |
| Cherry | Powdery mildew | 1052 |
| Coffee | Phoma | 6571 |
| Coffee | Cerscospora | 7681 |
| Coffee | Healthy | 18983 |
| Coffee | Rust | 8336 |
| Coffee | Miner | 16978 |
| Maize | Maize Lethal Necrosis | 3980 |
| Maize | Maize Streak Virus | 7217 |
| Maize | Cercospora leaf spot Gray leaf spot | 2390 |
| Maize | Abiotic | 77 |
| Maize | Rust | 2614 |
| Maize | Curvularia | 259 |
| Maize | Helminthosporiosis | 159 |
| Maize | Healthy | 8230 |
| Maize | Northern Leaf Blight | 3313 |
| Maize | Stripe | 2190 |
| Maize | Rust | 99 |
| Maize | Virosis | 148 |
| Cucumber | Healthy | 160 |
| Cucumber | Anthracnose | 160 |
| Cucumber | Bacterial Wilt | 160 |
| Cucumber | Downy Mildew | 160 |
| Cucumber | Gummy Stem Blight | 160 |
| Grape | Brown Spot | 782 |
| Grape | Downy Mildew | 392 |
| Grape | Mites | 400 |
| Grape | Black rot | 1834 |
| Grape | Esca (Black Measles) | 1383 |
| Grape | Healthy | 1329 |
| Grape | Leaf blight (Isariopsis Leaf Spot) | 1076 |
| Mango | Anthracnose | 500 |
| Mango | Bacterial Canker | 500 |
| Mango | Cutting Weevil | 150 |
| Mango | Die Back | 500 |
| Mango | Gall Midge | 500 |
| Mango | Healthy | 500 |
| Mango | Powdery Mildew | 500 |
| Mango | Sooty Mould | 500 |
| Orange | Haunglongbing (Citrus greening) | 5507 |
| Peach | Bacterial spot | 2297 |
| Peach | Healthy | 360 |
| Pepper | leaf spot | 67 |
| Black Pepper | Healthy | 1539 |
| Pepper | Bacterial spot | 997 |
| Black Pepper | yellow mottle virus | 273 |
| Black Pepper | leaf blight | 273 |
| Black Pepper | Healthy | 273 |
| Potato | Early blight | 1117 |
| Potato | Healthy | 152 |
| Potato | Late blight | 1105 |
| Raspberry | Healthy | 371 |
| Rice | Hispa | 1981 |
| Rice | Blast | 3005 |
| Rice | Bacterial Blight | 827 |
| Rice | Brown Spot | 2897 |
| Rice | Healthy | 3347 |
| Rice | Scald | 358 |
| Rice | Smut | 339 |
| Rice | Narrow Brown Spot | 353 |
| Soybean | Healthy | 5152 |
| Squash | Powdery mildew | 1964 |
| Strawberry | Healthy | 641 |
| Strawberry | scorch | 1109 |
| Sugarcane | Banded Chlorosis | 471 |
| Sugarcane | Brown Rust | 314 |
| Sugarcane | Brown Spot | 1722 |
| Sugarcane | Dried Leaves | 343 |
| Sugarcane | Grassy shoot | 346 |
| Sugarcane | Healthy | 430 |
| Sugarcane | Pokkah Boeng | 297 |
| Sugarcane | Sett Rot | 652 |
| Sugarcane | Smut | 316 |
| Sugarcane | mosaic virus | 663 |
| Sugarcane | Yellow Leaf | 1194 |
| Tea | Healthy | 1024 |
| Tea | Leaf blight | 496 |
| Tea | Red leaf spot | 500 |
| Tea | Red scab | 500 |
| Tomato | Verticulium wilt | 772 |
| Tomato | Bacterial floundering | 252 |
| Tomato | Fusarium | 18 |
| Tomato | Bacterial spot | 2226 |
| Tomato | Early blight | 2521 |
| Tomato | Healthy | 2918 |
| Tomato | Late blight | 2024 |
| Tomato | Leaf Mold | 1239 |
| Tomato | Septoria leaf spot | 4669 |
| Tomato | Spider mites Two-spotted spider mite | 1692 |
| Tomato | Target Spot | 1433 |
| Tomato | mosaic virus | 1173 |
| Tomato | Yellow Leaf Curl Virus | 5943 |
| Tomato | Mite | 390 |
| Tomato | Virosis | 1538 |
| Black gram | Cercospora leaf spot | 598 |
| Black gram | Healthy | 545 |
| Black gram | Insect | 408 |
| Black gram | Leaf Crinkle | 806 |
| Black gram | Yellow Mosaic | 1681 |
| Chili | Healthy | 2195 |
| Chili | Cercospora | 2217 |
| Chili | mites and trips | 2504 |
| Chili | nutritional | 2029 |
| Chili | powdery mildew | 2029 |
| Durian | Algal Leaf Spot | 105 |
| Durian | Healthy | 105 |
| Durian | Leaf blight | 103 |
| Durian | Leaf Spot | 100 |
| Eggplant | Healthy | 1451 |
| Eggplant | Insect | 546 |
| Eggplant | Leaf Spot | 602 |
| Eggplant | Mosaic Virus | 1362 |
| Eggplant | White Mold | 63 |
| Eggplant | Wilt | 65 |
| Guava | Anthracnose | 237 |
| Guava | Canker | 192 |
| Guava | Dot | 219 |
| Guava | Healthy | 1248 |
| Guava | Rust | 167 |
| Onion | Alternaria | 515 |
| Onion | Fusarium | 738 |
| Onion | Iris yellow virus | 282 |
| Onion | Healthy | 1105 |
| Onion | Purple blotch | 18 |
| Onion | Stemphylium leaf blight and collectrichum leaf blight | 90 |
| Onion | Virosis | 203 |
| Radish | Black leaf spot | 526 |
| Radish | Healthy | 613 |
| Radish | Mosaic Virus | 548 |
| Radish | Downey mildew | 601 |
| Radish | flea beete | 513 |
| Spinach | Bacterial Spot | 752 |
| Spinach | Healthy | 1399 |
| Spinach | Anthracnose | 102 |
| Spinach | Downey mildew | 240 |
| Spinach | Pest Damage | 513 |
| Soursop | Cutting Caterpillar | 631 |
| Soursop | Healthy | 656 |
| Soursop | Cutting Weevil | 642 |
| Soursop | Dieback | 652 |
| Soursop | Whiteflies | 636 |
| Soursop | Yellow | 621 |
| Wheat | Healthy | 102 |
| Wheat | Septoria | 97 |
| Wheat | Stripe rust | 208 |
| Hog_Plum | Healthy | 2829 |
| Hog_Plum | Insect | 953 |
| Bean | Blight | 510 |
| Bean | Healthy | 554 |
| Bean | Mosaic Virus | 562 |
| Bean | Rust | 568 |
| Cowpea | Bacterial wilt | 581 |
| Cowpea | Healthy | 536 |
| Cowpea | Mosaic virus | 579 |
| Cowpea | Septoria leaf spot | 577 |
| papaya | Anthracnose | 355 |
| papaya | Curl | 585 |
| papaya | Healthy | 228 |
| papaya | Bacterial Spot | 458 |
| papaya | Ring spot | 533 |
| watermelon | Downey mildew | 380 |
| watermelon | Mosaic Virus | 415 |
| watermelon | Healthy | 205 |

**Table S2 | Overview of image datasets used in this study.** *The table summarizes the original LeafNet data sources, followed by 13 additional datasets incorporated in this work (highlighted in bold). For each dataset, we report data provenance, geographic coverage, image acquisition conditions, background characteristics, crop species, imaging devices, and corresponding references.*

| **Dataset** | **Source type** | **Location** | **Image conditions** | **Background characteristics** | **Plant** | **Num. of raw images** | **Device** | **Ref.** |
| --- | --- | --- | --- | --- | --- | --- | --- | --- |
| PlantVillage | open-access | Multiple regions | Laboratory | Controlled | Multiple plant | 54304 | Digital camera | Hughes & Salathe, 2015 |
| JMuBEN2 | open-access | Kenya | Field | Natural | Coffee | 58555 | Digital camera | Jepkoech et al., 2021 |
| PlantDoc | open-access | Multiple regions | Field | Natural | Multiple plants | 2598 | Crowd-sourced | Singh et al., 2020 |
| CCMT Dataset | open-access | Ghana | Laboratory & Field | Controlled & Natural | Multiple plants | 24881 | Digital camera | Kwabena, 2023 |
| Grape Leaf dataset | open-access | Unknown | Field | Natural | Grape | 3000 | Digital camera | Anan, 2024 |
| Sugarcane Leaf dataset | open-access | India | Field | Natural | Sugarcane | 6748 | Mobile phone | Thite et al., 2024 |
| MangoLeafBD | open-access | Bangla- desh | Laboratory | Controlled | Mango | 1800 | Mobile phone | Ahmed et al., 2023 |
| Black Pepper Leaf Blight and Yellow Mottle Virus | open-access | Unknown | Laboratory | Controlled | Pepper | 273 | Unkno-wn | Melan, n.d. |
| Tea Leaf Dataset | open-access | Bangla-  desh | Field | Natural | Tea | 3960 | Mobile phone | Ahmad, 2024 |
| Maize Leaf Dataset | open-access | Tanzania | Field | Natural | Maize | 18148 | Mobile phone | Mduma & Laizer, 2023 |
| Cucumber | open-access | Bangla-  desh | Field | Natural | Cucumber | 1280 | Digital camera | Sultana et al., 2023 |
| Rice dataset | Research partner | Vietnam | Laboratory | Controlled | Rice | 13186 | Mobile phone | Quach et al., 2022 |
| Hog Plum | open-access | Bangla-  desh | Field | Natural | Hog Plum | 3782 | Mobile phone | Durjoy et al., 2025 |
| TOM2024 | open-access | Burkina Faso | Field | Natural | Tomato, maize & onion | 25844 | Digital camera | Appiah et al., 2025 |
| Beans and Cowpeas leaf dataset | open-access | Bangla-  desh | Field | Natural | Beans & Cowpea | 4467 | Mobile phone | Hasan et al., 2024 |
| BDPapaya-Leaf | open-access | Bangla-  desh | Field | Natural | Papaya | 2159 | Mobile phone | Mustofa et al., 2024 |
| Plant Pathology 2021 - FGVC8 | open-access | USA | Field | Natural | Apple | 18632 | Digital camera | Thapa et al., 2021 |
| Wheat dataset | open-access | Ethiopia | Field | Natural | Wheat | 407 | Unkno-wn | Getch, n.d. |
| Durian dataset | open-access | Vietnam | Field | Natural | Durian | 420 | Mobile phone | *Durian Diseases Dataset*, n.d. |
| Chilli dataset | open-access | India | Field | Natural | Chilli | 10987 | Unkno-wn | P, 2024a |
| Onion | open-access | India | Field | Natural | Onion | 4502 | Mobile phone | P, 2024b |
| SoursopBD | open-access | Bangla-  desh | Field | Natural | Soursop | 3838 | Mobile phone | Mustofa, 2025 |
| Radish | open-access | Bangla-  desh | Laboratory | Controlled | Radish | 2801 | Mobile phone | Hasan, 2025 |
| IDBGL | open-access | Bangla- desh | Laboratory | Controlled | Black gram | 4038 | Mobile phone | Shoib et al., 2025 |
| Eggplant | open-access | Bangla- desh | Laboratory | Controlled | Eggplant | 4089 | Mobile phone | Howlader et al., 2025 |
| Guava | open-access | Bangla- desh | Field | Natural | Guava | 3432 | Mobile phone | Shihab et al., 2025 |

**Table S3** | Overview of the task formulations supported in the LeafBench, including task abbreviations and corresponding descriptions.

| **Task Formulation** | **Abb.** | **Description** |
| --- | --- | --- |
| Disease Identification | DI | The core task of identifying the specific common name of the disease. It requires the model to distinguish between visually similar conditions across various crops (e.g., distinguishing between "Apple Scab" and "Apple Black Rot") |
| Pathogen Classification | PC | This goes beyond the disease name to identify the causal agent. The model must classify the primary cause of the infection—whether it is fungal, bacterial, viral, or a nutrient deficiency (abiotic) |
| Crop Species Identification | CSI | A foundational vision task where the model identifies the host plant (e.g., Apple, Corn, Potato). This is often a "sanity check" metric, as identifying the crop is a prerequisite for accurate disease diagnosis. |
| Symptom Identification | SI | Focuses on visual descriptors and fine-grained manifestations. Instead of naming the disease, the model describes the physical appearance—such as chlorosis (yellowing), necrosis (dead tissue), or specific lesion shapes (e.g., circular vs. angular). |
| Healthy-Diseased Classification | HDC | A binary classification task. This is the simplest level of reasoning, requiring the model to determine whether a leaf is entirely healthy or exhibits any pathological  symptoms. Most models perform exceptionally well (90%+) on this task. |
| Scientific Name Classification | SNC | This is a high-level taxonomic task. It requires the model to provide the formal binomial scientific name of the pathogen or the specific disease taxonomy (e.g., identifying that "Early Blight" in tomatoes is caused by the fungus Alternaria solani). |
| Lesion Identification | LI | The task of identifying individual lesion points or areas of infection. While classification looks at the whole leaf, LI evaluates the model's ability to focus on and identify specific "wounds" or spots on the leaf surface. |
| Leaf Symptom Detection | LSD | This involves spatial reasoning and detection. It tests if the model can accurately detect symptoms within the complex visual context of a leaf, often distinguishing the diseased area from shadows, dirt, or natural leaf variations. |
| Disease Severity Classification | DSC | An ordinal assessment of how far the disease has progressed. The model categorizes the  severity of the infection—usually into stages such as ”Early” and “Late”—based on the  percentage of the leaf area affected by symptoms. |

**Table S4 |** Representative examples of multiple-choice questions used for different task settings in LeafNet 2.0, showing the image input, question format, answer choices, and corresponding ground-truth labels.

| **Question Type** | **Image Name** | **Question** | **Options (A, B, C, D)** | **Answer** |
| --- | --- | --- | --- | --- |
| **DI** | Apple_Black_rot_0156.JPG | What disease is shown on the Apple leaf in this image? | A: Black rot, B: Powdery mildew, C: Narrow Brown Spot, D: leaf blight | **A** |
| **PC**( | Apple_Black_rot_0156.JPG | What type of pathogen is responsible for the disease shown in this image? | A: Fungal, B: Fungal, Bacterial, C: Virus, D: temperatures | **A** |
| **CSI** | Apple_Black_rot_0156.JPG | Which crop species does this diseased leaf belong to? | A: Apple, B: Pepper, C: Cassava, D: Black Pepper | **A** |
| **SI** | Apple_Black_rot_0156.JPG | Which of the following best describes the main symptoms visible on the leaf? | A: Small black spots, concentric rings, B: tan-brown lesion..., C: brown lesions..., D: irregular chlorotic areas | **A** |
| **HDC** | Apple_Black_rot_0156.JPG | How would you classify the leaf shown in this image? | A: Healthy, B: Diseased | **B** |
| **SNC** | Apple_Black_rot_0156.JPG | What is the scientific name of the pathogen causing the disease shown in this image? | A: Botryosphaeria spp., B: Gymnosporangium juniperi-virginianae, C: Elsinoë mattei, D: Colletotrichum falcatum | **A** |
| **LI** | Apple_Black_rot_0156.JPG | Which of the following best describes the lesion characteristics visible on the leaf in this image? | A: small, reddish- to purplish-brown specks..., B: Small bluish spots..., C: Small, circular, greenish-yellow spots..., D: Yellowish, oily or angular... | **A** |
| **LSD** | Apple_Black_rot_0156.JPG | What are the primary symptoms of the apple leaf infected with Black rot? | A: Brown to dark necrotic lesions with darker margins, B: Yellowing of the entire leaf, C: Small white spots..., D: Curling of the leaf edges | **A** |
| **DSC** | Apple_Black_rot_0156.JPG | What stage of disease progression is visible in the image? | A: Late, B: Early | **A** |
